# Steady-State Visually Evoked Potentials as Readouts for Abnormal GSK3β Activity in *Drosophila*

**DOI:** 10.1101/2025.10.24.684304

**Authors:** Oscar Solis, Toby Shaw-McGrath, Alex R. Wade, Ines Hahn

## Abstract

Glycogen synthase kinase 3β (GSK3β) is important in neuronal development and maintenance, acting as a key regulator by controlling a broad range of cellular processes. The effects of its dysregulation range from impairments in axonal transport and energy metabolism to synapse formation and neuronal plasticity. However, how such cellular defects link to neuronal dysfunction is less well studied despite the links between GSK3β and various neurological conditions, such as Alzheimer’s and Parkinson’s diseases, mood disorders and autism. Here we used a steady-state visually evoked potential (SSVEP) assay with *Drosophila* to measure the effect of altering GSK3β expression on neuronal function. We recorded SSVEPs from flies that expressed constitutively active or inactive (kinase-dead) variants of GSK3β and showed that the visual responses of these flies differed from those of controls. This was indicated by increased response latency and reduced photoreceptor maximum response (*R*_*max*_). Importantly, multivariate pattern classification can distinguish between over- and inactive GSK3β conditions. Taken together, we show that SSVEPs provide a powerful tool for future studies to link the molecular and cellular functions of GSK3β with their effects on neuronal function. Furthermore, we introduce a monitoring protocol that ensures the quality of datasets to support future fly SSVEP experiments.

## Introduction

Within the nervous system, glycogen synthase kinase 3β (GSK3β) is required for almost all stages of neuronal development and maintenance (Hur & Zhou, 2010). It also plays a key role in neuronal plasticity (Jaworski et al., 2019). Overactive GSK3β is associated with many neurodegenerative disorders, including Parkinson’s (Duka et al., 2009; Nagao & Hayashi, 2009) and Alzheimer’s (Pei et al., 1997; Uemura et al., 2007) diseases. GSK3β is also related to neurodevelopmental conditions such as autism spectrum disorder (Rizk et al., 2021) and psychiatric disorders such as bipolar disorder (Chatterjee & Beaulieu, 2022; Jope & Roh, 2006), Targeting GSK3β has seen the most success as a treatment for psychiatric disorders, although this comes with some limitations (Benedetti et al., 2013; Gitlin, 2016). For neurodegenerative disorders, targeting GSK3β has shown promise in animal models (Li et al., 2020; Youdim & Arraf, 2004) but often fails to pass human clinical trials (Hampel et al., 2009; Lovestone et al., 2015; Ser et al., 2013).

GSK3β is a multifaceted kinase with many targets (>100) (Eleonore Beurel et al., 2015). These include metabolic proteins such as transcription factors, the machinery involved in proteostasis, and structural proteins such as microtubule binding proteins (Hajka et al., 2021). In neurons, GSK3β is related to dendritic and axonal outgrowth (Garrido et al., 2007; Gärtner et al., 2006; Jiang et al., 2005; Owen & Gordon-Weeks, 2003; Rui et al., 2013) as well as axonal maintenance (Voelzmann et al., 2025). More broadly, GSK3β has key roles in healthy physiological processes such as in metabolism (L. Wang et al., 2022) and apoptosis (Eléonore Beurel & Jope, 2006). Thus far, much of the focus has been on understanding the roles of GSK3β at the molecular and cellular levels.

To bridge this understanding to neuronal dysfunction at the systems level, we propose the use of steady-state visually evoked potentials (SSVEPs) recorded from *Drosophila melanogaster*. SSVEPs are population electrophysiological responses to a periodic visual stimulus (Norcia et al., 2015; Wade & Baker, 2025). The use of the technique in flies to assay disease-related genotypes was developed by Afsari et al. (2014) who found that flies with Parkinson’s disease-related mutations in the *LRRK2* gene showed abnormal visual gain control compared to control flies. Using this technique in a fly model of autism, Vilidaite et al. (2018) also showed that the pattern of deficits across developmental stages can be strikingly similar between flies and humans. The main advantage of the technique lies in the high signal-to-noise ratios that come from analysing the data in the frequency domain and discarding information at frequencies unrelated to the stimulus (Norcia et al., 2015). In addition, population responses from different stages of the fly visual system – namely the photoreceptors, lamina neurons and medulla neurons – can be isolated (Afsari et al., 2014) and often add important information about the different visual processing stages (Himmelberg et al., 2018).

Here we present data and the results of statistical analyses that demonstrate how SSVEPs can be used as a readout for abnormal GSK3β activity in flies. We compared recordings from control flies with flies expressing either constitutively active (overactive) or kinase-dead (inactive) forms of *Drosophila* GSK3β (shaggy) in neurons. We analysed the SSVEP data in three ways: as done classically by modelling contrast-response functions (CRFs) (Afsari et al., 2014), incorporating phase (Baker, 2021), and using multivariate pattern analyses (Himmelberg et al., 2018; West et al., 2015). We find that differences in CRFs are best described by a reduction in response gain control in the photoreceptors of flies with abnormal GSK3β activity, and that these flies also have slightly altered response latency (phase). Despite similar phenotypes observed in flies expressing overactive and inactive GSK3β on neuronal level (Voelzmann et al., 2025), a pairwise classifier was able to distinguish between the two variants with a mean accuracy of 76%. We discuss the potential of this experimental model in further investigating the links between the cellular and molecular roles of GSK3β to neuronal function at the systems level.

## Materials and methods

### Drosophila stocks, genetics and maintenance

The GAL4-UAS system (Brand & Perrimon, 1993) was used to target the overexpression of mutants of *shaggy* (*sgg*), the fly orthologue for GSK3β, pan-neuronally (Figure 1A). Male flies with the UAS constructs *UAS-sgg*^*S9A*^ (constitutively active) or *UAS-sgg*^*A81T*^ (kinase dead; both 3^rd^ chromosomal; Bloomington stocks #5362 and #5360 respectively) (Bourouis, 2002) were crossed with female *elav-GAL4* flies (3^rd^ chromosomal; Bloomington stock #8760) (Luo et al., 1994). From the F_1_ generation, male *elav-GAL4/ UAS-sgg*^*S9A*^ (hereafter ‘GSK3β^CA^’) or *elav-GAL4/ UAS-sgg*^*A81T*^ (hereafter ‘GSK3β^DN^’) flies were aged 7 days post-eclosion prior to testing. 7-day old male *elav-GAL4* flies were used as controls. All fly lines were raised in vials containing standard media, at a constant temperature of 25°C, and in a 12hr:12hr light/dark cycle. Data were collected from 20 flies for each genotype.

**Figure 1.**
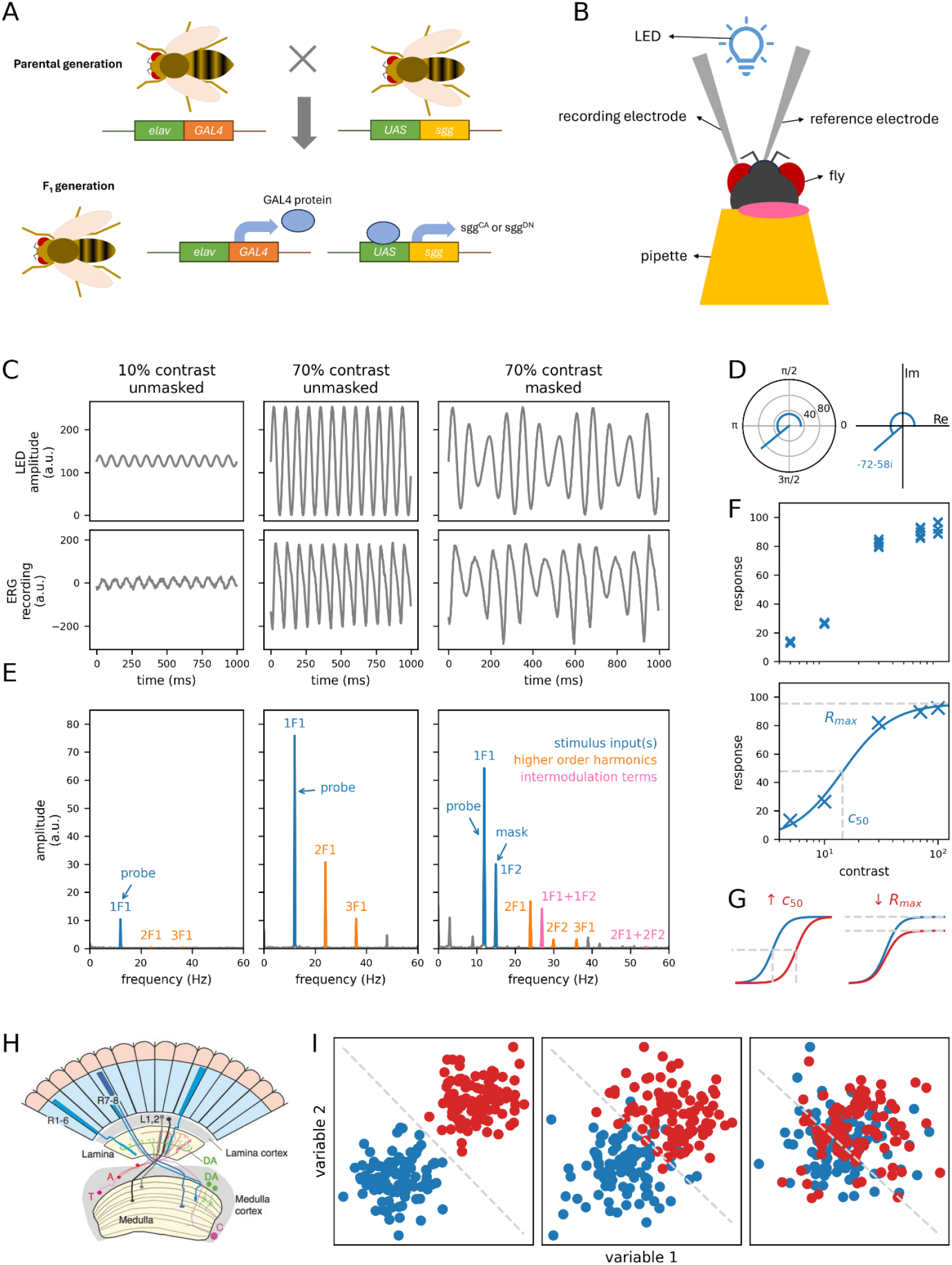
Experimental pipeline consisting of recording, preprocessing and analysis of steady-state visually evoked potentials (SSVEPs). **(A)** The GAL4-UAS system was used to induce over-expression of in/active *shaggy* (*sgg*) mutants in neurons. **(B)** A fly is mounted into the electrophysiology rig and positioned directly in front of the blue LED which presents the stimulus. **(C)** Representative stimuli (top) and recordings (bottom) for three stimulus conditions. **(D)** The outputs of a Fourier transform are complex numbers which can be plotted in polar space (left plot) with constituent amplitude (line from origin) and phase (curve depicting angle) or Cartesian space (right plot) with real (Re) and imaginary (Im) parts. **(E)** Representative amplitude spectra for the same conditions as in (C). **(F)** An exemplar contrast-response function (CRF) from an individual fly (top) which is modelled by a three-parameter hyperbolic function (bottom). Small crosses indicate the amplitude of the response at a given frequency (here, 12Hz) for each trial. Large crosses are coherently averaged amplitudes used to model the hyperbolic function shown as the blue curve. **(G)** Changes in *c*_50_ correspond to vertical scaling (left) whereas changes in *R*_*max*_ correspond a horizontal scaling of the hyperbolic curve. **(H)** Stages of the fly visual system, such as the photoreceptors, lamina neurons and medulla neurons, correspond to components in the SSVEP amplitude spectrum. Figure from Afsari et al. (2014). **(I)** Simulated 2-dimensional feature spaces showing how a linear discriminant analysis (LDA) can be used as a classification technique to distinguish between two classes (here, ‘blue’ and ‘red’). Each dot on these plots is a ‘fly’. Based on features ‘variable 1’ and ‘variable 2’, the data may form distinct clusters (left panel) that can be separated by a single linear boundary (grey line). The class of a new fly can be determined based on which side of the boundary the fly is in this feature space. It is more likely that these clusters overlap (middle panel), reducing the accuracy of the classification. It is also possible for these clusters to overlap significantly (right panel), resulting in a classifier that performs at around chance level (50% with two classes).

### Steady-state visually evoked potential (SSVEP) assay

Flies were aspirated into a shortened pipette using a pooter such that its head protruded through (Figure 1B). Nail varnish was applied behind the neck to mount the flies onto the pipette tip. A combined stimulus display/recording system, which consisted of a set of high-intensity LEDs modulated by pulse-width modulation and high-impedance amplifier controlled by a dedicated microcontroller (Arduino Due), generated and presented all stimuli and recorded electrophysical responses. Flies were mounted into this system such that it was directly in front of the LEDs. Electrodes filled with saline solution (130 mM NaCl, 4.7 mM KCl, 1.9 mM CaCl_2_) (West et al., 2015) were gently placed on the eye and mouthpart.

The contact between the electrodes and the fly eyes was tested through an electroretinography protocol that consisted of 5 pulses of blue light. Traces generated online during recording were visually inspected to confirm contact. The setup was adjusted once if required and flies were discarded after 2 protocols were run.

The main SSVEP protocol consisted of blue light flickering at 12Hz for 4000ms at contrasts of 5%, 10%, 30%, 70% and 100%. In some trials, an additional 15Hz masking stimulus was added to the 12Hz stimulation frequency (except for 100% contrast trials). This resulted in a total of 9 unique stimulus conditions repeated 5 times each for a total of 45 trials for each fly. Exemplar stimuli and recordings for representative stimulus conditions are shown in Figure 1C.

Traces were visually inspected both online and offline to monitor the quality of the data. One-sample t-tests were implemented to test whether responses for each stimulus condition were significantly above noise (see Figure 2B). A conservative criterion of p <.001 for all stimulus conditions was required for fly data to qualify for data analysis.

**Figure 2.**
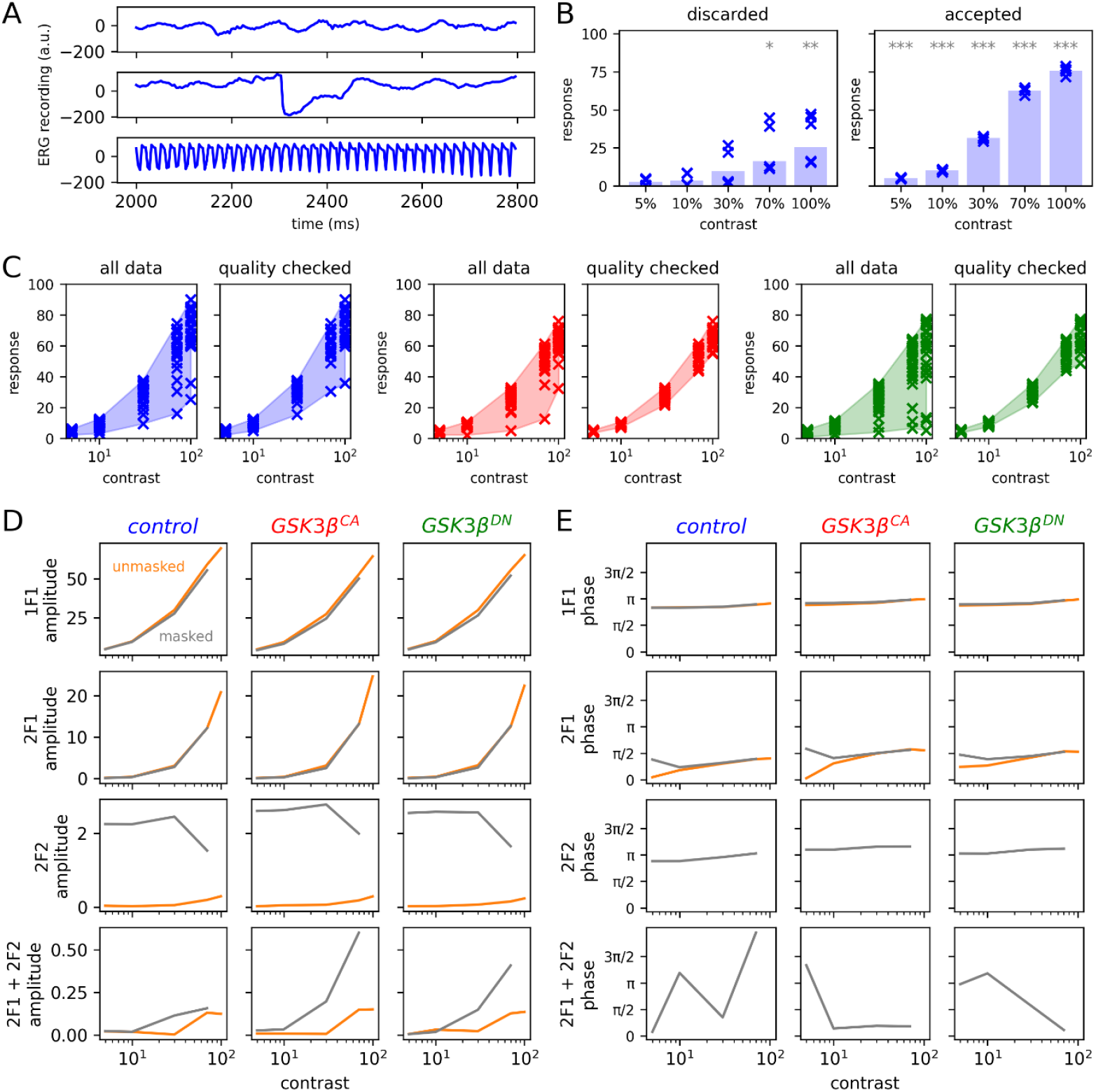
Quality control procedures improve precision of raw data. The data plotted in (A) and (B) are unmasked responses from control flies. **(A)** Representative traces showing an intact recording (top trace), a recording affected by movement (middle trace) and a noisy, aberrant recording (bottom trace) for 5% contrast. **(B)** Data from individual flies were discarded from or accepted for further analysis based on whether (right graph) or not (left graph) the statistical significance of a one-sample t-test of the amplitudes for a given contrast was less than.001 (indicated by ***). Crosses represent amplitudes from individual trials and bars represent their mean. **(C)** Coherently averaged responses from flies, with each fly represented by a cross, for all samples (left graphs) and after the removal of discarded flies (right graphs, n = 20) for control (blue), GSK3β^CA^ (red) and GSK3β^DN^ (green) flies. Shaded areas represent the range of the data. **(D, E)** Grand means of coherently averaged mean amplitudes (D) and mean phases (E) plotted against contrast for unmasked (orange) and masked (grey) stimuli.

Data preprocessing steps, analyses and visualisation were implemented using Python 3.10. A Fast Fourier Transform (Cooley & Tukey, 1965) was used to transform the data into the frequency domain, the outputs for which are complex numbers which can be identified by their amplitude and phase (Figure 1D, left) or their real and imaginary parts (Figure 1D, right). Apart from peaks at the stimulation frequency of 12 Hz, amplitude-frequency spectra also contain peaks related to this frequency either as harmonics or intermodulation components (Figure 1E).

### Modelling contrast-response functions (CRFs)

For each fly, amplitudes for each stimulus condition were calculated after coherently averaging across trials. These amplitude data were then fit to a hyperbolic ratio function (Figure 1F) (Albrecht & Hamilton, 1982) as in previous work from our lab (Afsari et al., 2014; Tsai et al., 2012):

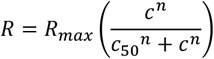

where *R* and *c* represent response amplitudes and contrasts respectively, *R*_*max*_ represents the modelled maximum response of the neuronal population compared to baseline, *c*_50_ represents the semisaturation contrast at which the response is half of *R*_*max*_, and the exponent *n* determines the steepness of the non-linear region of the curve and represents the sensitivity of the neuronal population to dynamic changes in the stimulus. We kept the value of *n* = 2 in line with previous studies in both flies and humans (Afsari et al., 2014; Tsai et al., 2012). The effects of changes in the values of *c*_50_ and *R*_*max*_ to the curve is shown in Figure 1G. Responses at different harmonics of the input frequency correspond to activity at different layers of the fly visual system (Figure 1H) (Afsari et al., 2014). We use this model for 1F1 and 2F1 components which reflect activity from the photoreceptors and lamina neurons.

### Multivariate Classification

We used linear discriminant analysis (LDA) as a classification technique (Figure 1I) and assessed its performance via a ‘leave-one-out’ (LOO) analysis as done previously in our lab (West et al., 2015; Himmelberg et al., 2018). We performed pairwise classifications based on 72 features: 9 unique stimulus conditions × 4 representative frequencies (1F1, 2F1, 2F2 and 2F1 + 2F2) × 2 values to identify complex values in Cartesian space (real and imaginary parts). The LOO analysis involved training the classifier on data pooled from all except one fly and then predicting the class of the ‘unseen’ fly. This is repeated over all flies to obtain a classification accuracy.

### Statistical analysis

Bootstrapping (Efron & Tibshirani, 1993) was used to analyse whether differences in the parameters of modelled CRFs between genotypes and the performance of the classifier compared to chance levels (50%) were statistically significant. The data were resampled with replacement to generate 1000 ‘synthetic’ datasets which were used to either fit the hyperbolic ratio function to estimate *R*_*max*_ and *c*_50_ or to calculate the accuracy of a classifier assessed using LOO. For differences in CRF parameters, a two-sided p-value was calculated as double the lesser fraction of bootstraps for control flies at either side of the bootstrapped mean of a GSK3β variant. For classification accuracies, a p-value was calculated as the proportion of bootstrapped accuracies less than the chance level of 50% (West et al., 2015).

To take phase information into account, we compared visual response with the bivariate 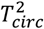 statistic (Victor & Mast, 1991) based on R code from Baker (2021).

## Results

### Data quality checks improve the precision of modelling

As a first step, we aimed to introduce quality control measures to ensure that data is suitable for subsequent modelling and analysis, allowing us to detect even subtle changes in vision between conditions. The quality of the data can be affected by how well the fly was mounted throughout recording (Figure 2A). Although traces were inspected visually as they were generated, loss of contact between the electrodes and fly eye may sometimes be missed. This leads to the formation of clustered responses of high and low amplitudes when the data are plotted on a CRF (Figure 2B, left). To determine the quality of the data objectively, and because only SSVEP *amplitudes*, as opposed to taking phase into account, are classically analysed at this stage (Afsari et al., 2014), one-sample t-tests were conducted on amplitude data for individual flies and a strict level of significance was employed (α =.001) (Figure 2B, right). Data from flies that did not meet this criterion were omitted from further analysis and replaced until a sample size of 20 flies for each genotype was reached. The result of this exclusion criteria on the dataset and modelling can be seen in Figures 2C-E for each genotype.

### Abnormal GSK3β activity reduces photoreceptor response gain control

To evaluate whether abnormal GSK3β activity leads to systems-level neuronal dysfunction, we expressed constitutively active (GSK-3β^CA^) or inactive GSK-3β (dominant negative, GSK-3β^DN^) mutants in neurons using *elav-GAL4* and recorded SSVEPs from the fly eye. *elav-GAL4*-only flies served as controls. Here we analysed these recordings by modelling the amplitudes of responses as CRFs with two parameters: *R*_*max*_ representing the maximum response of the neuronal population and *c*_50_ representing the semi-saturation constant, which can be interpreted as the sensitivity of the neuronal population to contrast. We model the 1F1 and 2F1 components as previous work has shown that these correspond to photoreceptor and lamina neuron activity respectively (Afsari et al., 2014) allowing us to see the effects of abnormal GSK3β activity at different stages of the fly visual system.

The results of the bootstrapping analyses are shown in Figures 3A and 3B and the fitted metrics are shown in Figures 3C and 3D. For the 1F1 component, GSK3β^DN^ expressing flies had lower *c*_50_ for unmasked stimuli than in control flies (*p* =.042); this effect was not observed with masking. *R*_*max*_ for unmasked stimuli is reduced in both GSK3β^DN^ (*p* =.002) and GSK3β^CA^ (*p* <.001) flies compared to control flies. *R*_*max*_ for masked stimuli is also reduced in GSK3β^DN^ (*p* =.028) and GSK3β^CA^ (*p* =.012) flies compared to control flies. For the 2F1 component, *c*_50_ was greater in GSK3β^CA^ flies compared to control flies (both unmasked and masked), though this increase was only significant in response to masked stimuli (*p* =.008). There were no effects of GSK3β activity on 2F1 *R*_*max*_.

**Figure 3.**
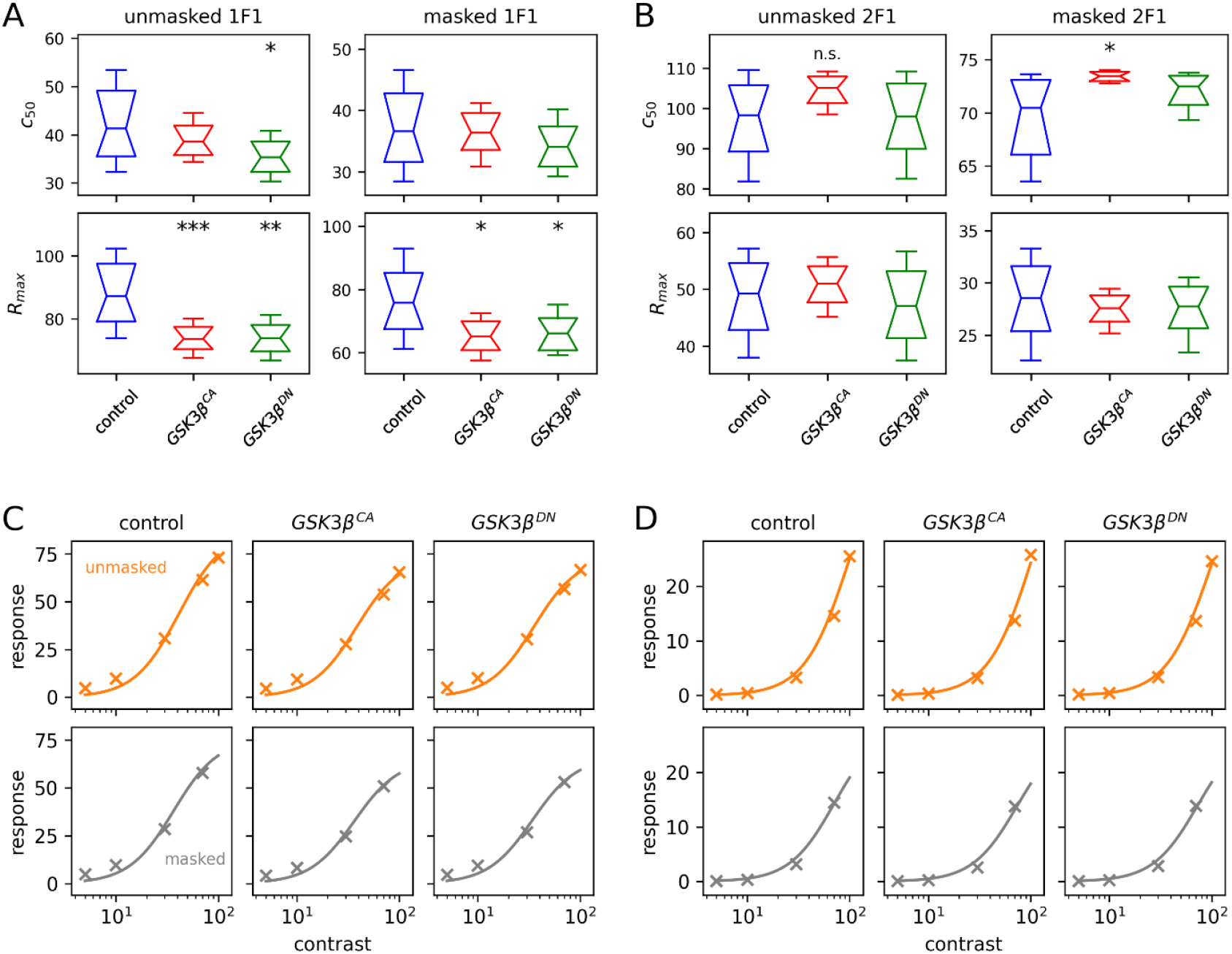
Modelling contrast-response functions (CRFs) reveals differences between control flies and flies with abnormal GSK3β activity. **(A, B)** Box plots showing the means of bootstrap estimates of the parameters, *c*_50_ (top) and *R*_*max*_ (bottom), for unmasked (left) and masked (right) responses from 1F1 (A) and 2F1 (B) steady-state visually evoked potential (SSVEP) components. Notches and whiskers indicate the 95% confidence interval and range of bootstrap estimates respectively. Significant differences in comparison to control flies are indicated by asterisks: * = *p* <.050; ** = *p* <.010, *** = *p* <.001. ‘n.s.’ indicates a non-significant difference. **(C, D)** Hyperbolic functions modelling CRFs based on the means of bootstrapped parameters for 1F1 (C) and 2F1 (D) components for unmasked (top) and masked (bottom) responses. Scatterplots are the coherently-averaged mean amplitudes as shown in Figure 2 and are fitted well by the modelled CRFs.

Collectively, these results suggest that expression of either overactive or inactive GSK3β are associated with defects in the fly visual system that are best described as a reduction in response gain control in their photoreceptors. However, there are some differences between the two GSK3β variants which can be observed in their responses to unmasked stimuli: increased contrast sensitivity of the photoreceptors in GSK3β^DN^ flies and decreased contrast sensitivity of the lamina neurons in GSK3β^CA^ flies.

### Abnormal GSK3β activity affects amplitudes and phase

The Fourier transform of periodic signals generates complex numbers that have both an amplitude and a phase. Whereas scalar amplitudes have classically been modelled as CRFs, phase is often omitted but is informative of the response latency of the signal. This temporal lag has been reported by the original *Drosophila* SSVEP study of Parkinson’s disease-related mutations in the LRRK2 protein (Afsari et al., 2014).

To include phase in the comparisons of visual responses between flies with overactive or inactive GSK3β to control flies, the data were analysed using the *T*_*circ*_^2^ statistic (Baker, 2021; Victor & Mast, 1991). Upon visual inspection, the difference in phase lag between control and flies with modified GSK-3β activity is qualitatively similar across contrasts and harmonic/intermodulation terms (Figures 4A to 4B). *T*_*circ*_^2^ statistical tests were conducted between control and abnormal GSK3β flies (Figure 4C). For responses at 100% contrast, there are significant differences between control and GSK3β^CA^ flies in the unmasked 1F1 (*T*_*circ*_^2^ = 0.782, *F*(2,76) = 7.819, *p* <.001) and 2F1 (*T*_*circ*_^2^ = 0.814, *F*(2,76) = 8.142, *p* <.001) components. Similarly, there are significant differences between control and GSK3β^DN^ flies in the unmasked 1F1 (*T*_*circ*_^2^ = 0.602, *F*(2,76) = 6.019, *p* =.004) and 2F1 (*T*_*circ*_^2^ = 0.388, *F*(2,76) = 3.884, *p* =.025) components. Examination of the 2F2 component, which is indicative of responses to the masking stimulus, also revealed differences at 30% contrast in responses for GSK3β^CA^ (*T*_*circ*_^2^ = 1.044, *F*(2,76) = 10.439, *p* <.001) or GSK3β^DN^ (*T*_*circ*_^2^ = 0.407, *F*(2,76) = 4.067, *p* =.021) flies compared to controls. Apart from at masked 70% contrast for GSK3β^CA^ flies, we observed no other statistically significant differences between genotypes for the 2F1 + 2F2 component, though this may be due to poor signal-to-noise ratio.

**Figure 4.**
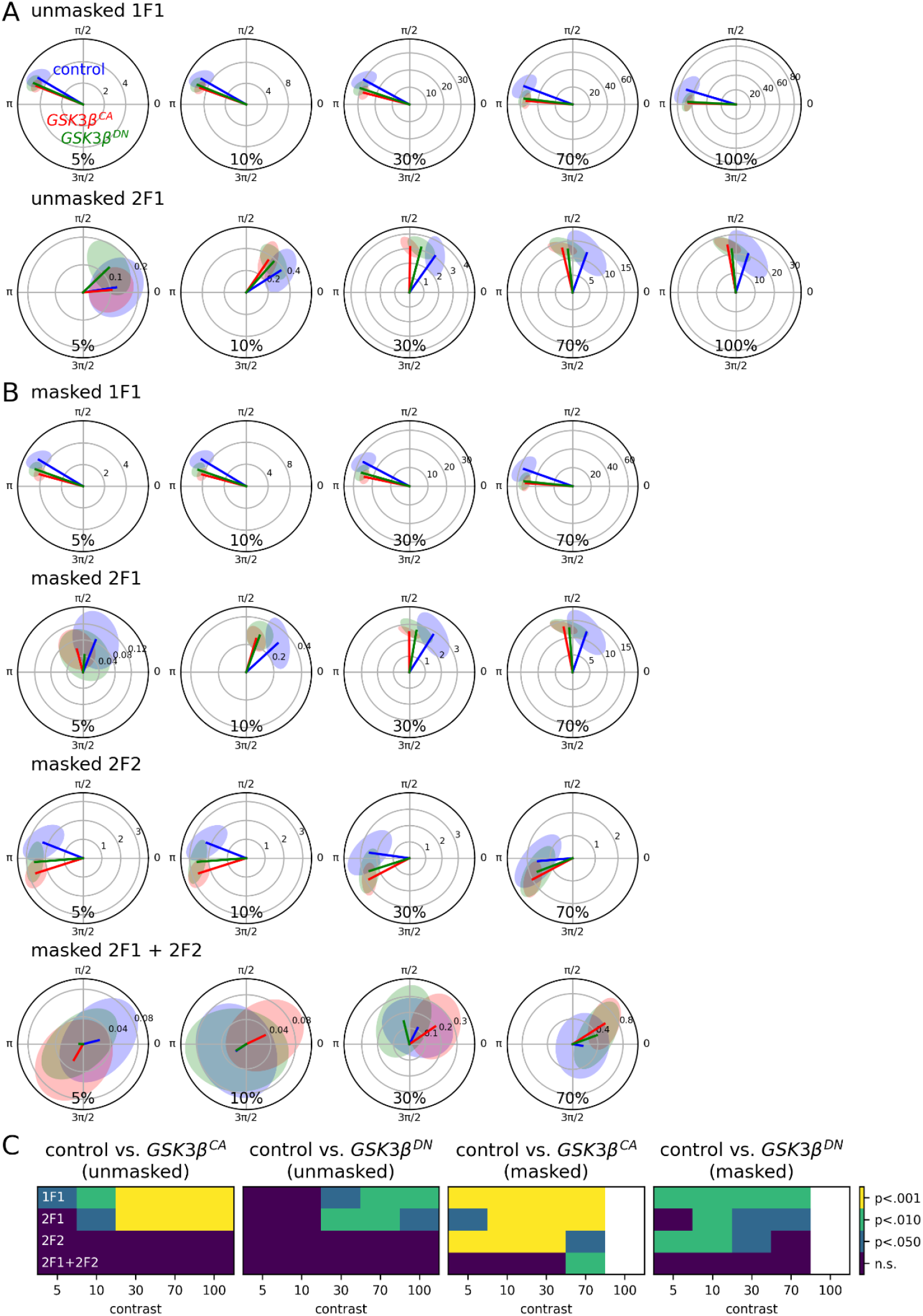
Statistical analysis including both amplitude and phase information reveals differences between control flies and flies with abnormal GSK3β activity. **(A, B)** Polar plots showing the mean amplitude (opaque line) and bounding error ellipse (shaded area) for unmasked (A) and masked (B) stimuli for each stimulus contrast for each genotype. Plots are shown for the stimulation frequency (12Hz labelled 1F1, first row) and the second harmonic (24Hz labelled 2F1, second row). For masked responses, additional plots are displayed for the second harmonic of the masking frequency (30Hz labelled 2F2, third row) and a fourth-order intermodulation term (54Hz labelled 2F1 + 2F2, fourth row). **(C)** Heatmaps showing uncorrected p-values from statistical analyses using the 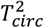 statistic: n.s. = non-significant.

These results suggest that GSK3β activity may also be related to the phase delay of neuronal populations, which is similarly perturbed when the kinase is over-or inactive.

### Pairwise classification distinguishes between GSK3β variants

As proof-of-concept, we also conducted multivariate analyses. Previously, this has been used to show that SSVEP responses can be used to classify a broad range of Parkinson’s-related genetic mutations and that the disease genotypes can also be distinguished from healthy genotypes (Himmelberg et al., 2018; West et al., 2015). Additionally, we are also able to use multivariate analyses to determine whether responses between GSK3β^CA^ and GSK3β^DN^ flies as this approach does not require data from one genotype as a null distribution.

Pairwise classifiers based on LDA were trained using features from four components (1F1, 2F1, 2F2, 2F1 + 2F2) which are representative of different stages of the fly visual system (Afsari et al., 2014). In addition to the 9 unique stimulus conditions and real and imaginary parts to each complex output, 72 features were used by the classifier to draw a linear boundary to separate two classes.

The accuracies of the classifiers assessed using LOO analysis coupled with bootstrapping is shown in Figure 5. Controls can be reliable distinguished from flies expressing GSK3β^CA^ (*p* <.001) and GSK3β^DN^ (*p* <.001) with accuracies of 95% and 87% respectively. The two GSK3β variants can also be classified from each other with an accuracy of 76% (*p* =.004). Finally, given that similar phenotypes have been observed in the previous analyses in terms of photoreceptor *R*_*max*_ and phase delay, we grouped both GSK3β^CA^ and GSK3β^DN^ flies into a single class (‘modified GSK3β’) and compared this against control flies. This classifier had an accuracy of 86% (*p* <.001).

**Figure 5.**
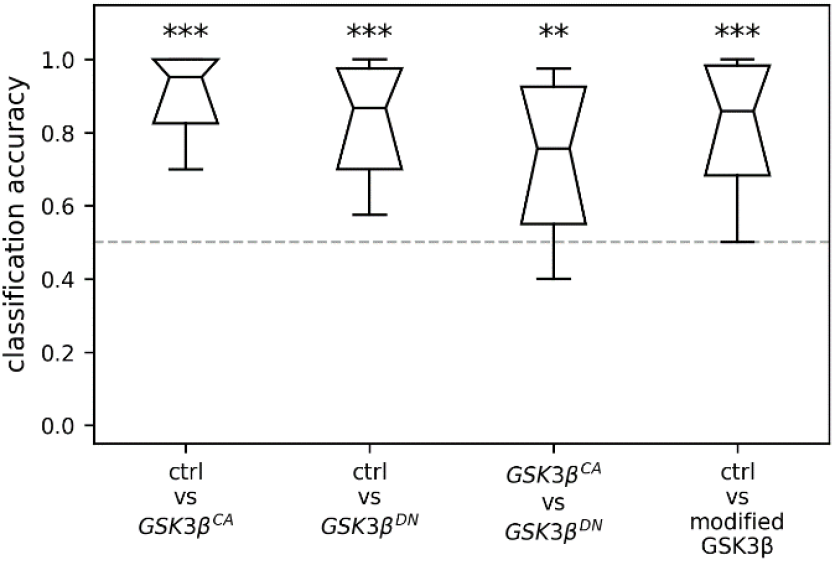
Linear discriminant analysis allows accurate pairwise classification of genotype based on steady-state visually evoked potentials (SSVEPs). Box plots show the means of bootstrap estimates of classification accuracies, with notches and whiskers indicating the 95% confidence interval and range of the bootstrap estimates respectively. The grey line indicates ‘chance’ performance of the classifier at 50%. Significant performance is indicated by asterisks: ** = *p* <.010, *** = *p* <.001.

These results are consistent with the previous analyses that showed that visual defects are associated with abnormal GSK3β activities. However, these results suggest that there are sufficient differences between over- and inactive GSK3β variants to be able to discriminate between them.

## Discussion

Here we have created a pipeline for fly SSVEP experiments that include quality control of the raw data and several approaches to data analysis. We used this pipeline to assess if the mis-regulation of GSK3β, a key driver of neurological disorders, affects electrophysiological responses in the fly visual system. We found evidence that flies expressing either constitutively active or kinase-dead, i.e., inactive, forms of GSK3β exhibited altered visual responses compared to controls. This was observed as reduced photoreceptor maximum response (Figure 3), best described as a reduction in response gain control, as well as increased response latency (Figure 4) indicated by temporal lags that were similar across photoreceptors and lamina neurons and even as a response to masking. Although *Drosophila* models of GSK3β appear to be showing similar defects – *R*_*max*_ and response latency but also cellular readouts such as microtubule organisation (Voelzmann et al., 2025) – multivariate pattern classification was able to distinguish between different genotypes (Figure 5). This suggests that different cellular and physiological mechanisms may be affected by dysregulated GSK3β.

The precision afforded by the data quality control measures can be observed in Figure 2. To further explore this, we repeated the experiment without these measures with the same genotypes – i.e. *elav-GAL4* driving the expression of overactive or inactive GSK3β – and analysed the data as modelled contrast response functions (Supplementary Figure 1). While we observed a comparable reduction in photoreceptor maximum response in the unmasked condition for flies expressing overactive or inactive GSK3β, there are noticeable differences for other parameters. It is possible that these differences result from using a different display/recording apparatus. However, given the qualitatively wider 95% confidence intervals, it is likely that the differences in effects are due to difference in precision. Nevertheless, the effect on photoreceptor *R*_*max*_ when modifying GSK3β activity appears to be robust between the parallel approaches.

Besides vision, GSK3β has also been investigated in behaviours such as habituation, the reduction of any response to frequently occurring stimuli. Over time, the stimulus is considered redundant and noninformative, so a heightened response would be unnecessary. Overactive GSK3β in flies results in greater levels of habituation compared to inactive GSK3β (Wolf et al., 2007). These findings, together with those in the present study, highlight the multifunctionality of GSK3β, with various roles not just within neuronal development but also in a range of other physiological processes.

The fact that both overactive and inactive GSK3β altered visual responses in flies suggests that GSK3β activity needs to be tightly balanced. Previous studies have indicated important roles for GSK3β in photoreceptor function and survival (Kisseleff et al., 2021; Sugie et al., 2015). There are a number of mechanisms by which GSK3β could affect function. For example, GSK3β was reported to negatively regulate pre-synaptic vesicle exocytosis by interfering with Ca2+-dependent SNARE complex formation in rat hippocampal neurons (Zhu et al., 2010). It controls synaptic plasticity by stabilising N-methyl-d-aspartic acid (NMDA) receptors at synapses (Amici et al., 2021). Additionally, active endogenous GSK3β is required to maintain excitatory AMPA receptors at the synaptic membrane (Wei et al., 2010).

Misregulation of GSK3β during development may also underlie the functional impairments observed in this study: active GSK3β seemingly promotes axon formation and prevents dendrite formation whereas inactive GSK3β upregulates dendritic growth and negatively affects axonal growth (Garrido et al., 2007; Gärtner et al., 2006; Jiang et al., 2005; Owen & Gordon-Weeks, 2003; Rui et al., 2013). In flies, activation of GSK3β *decreases* synaptic growth and plasticity at neuromuscular junctions (Franco et al., 2004) while loss of GSK3β has the opposite effect. Expressing overactive GSK3β or using GSK3β inhibitors in neuronal cultures leads to atypical dendritic and axonal morphology (Garrido et al., 2007; Gärtner et al., 2006; Jiang et al., 2005; Owen & Gordon-Weeks, 2003; Rui et al., 2013) and defects in microtubule networks (Voelzmann et al., 2025). Additionally, multiple neurodegenerative diseases are associated with overactive GSK3β (Nagao & Hayashi, 2009; Pei et al., 1997; C. Wang et al., 2023). However, trials using global GSK-3β inhibitors have mostly failed (Cheng et al., 2024) suggesting that its complete inhibition is neurotoxic. Defects observed with inactive GSK3β suggests that when developing drug treatments targeting GSK3β, the treatment must target the overactive GSK3β precisely to reduce the risk of affecting other physiological processes.

Taken together, the evidence suggests that proper neuronal function requires maintaining a precise equilibrium between the active and inactive states of GSK3β and that the balance of GSK3β activity must be tightly controlled.

The functions and molecular roles of GSK3β are diverse. Currently, our understanding of this important kinase is obstructed by our inability to systematically analyse the links between its molecular functions, its many cellular targets, the many processes it regulates and its impacts on neuronal physiology and function. We propose that combining *Drosophila* genetics with its detailed cellular readouts in our established primary culture model (Hahn et al., 2021; Qu et al., 2022, 2019, 2017) with fly SSVEP experiments to assess neuronal function is a powerful tool to bridge this gap in the future.

---

**Supplementary Figure 1.**
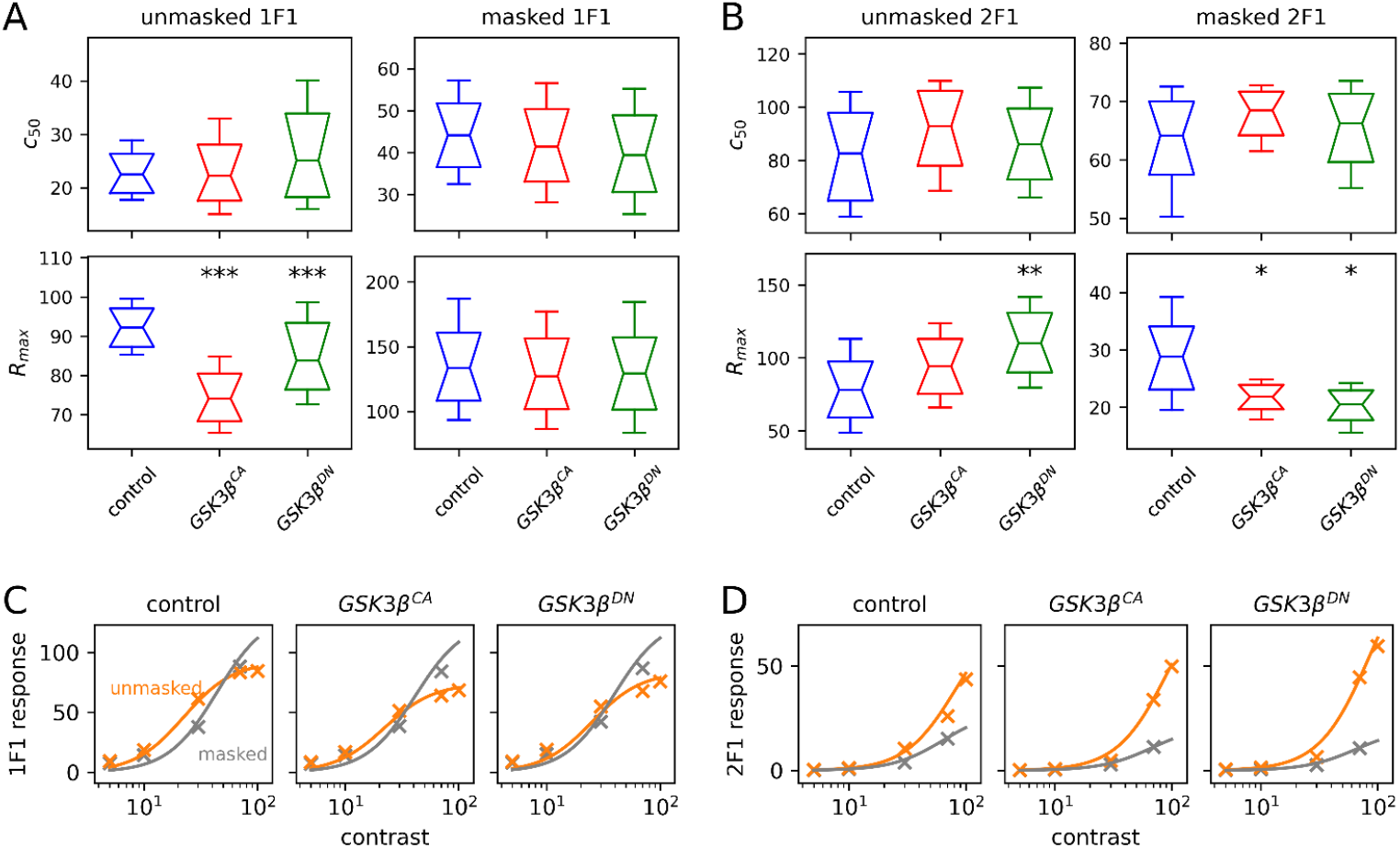
Modelling contrast-response functions (CRFs) from an independent repeat of the SSVEP experiment replicates decrease of photoreceptor *R*_*max*_ in flies expressing GSK3β variants. **(A, B)** Box plots showing the means of bootstrap estimates of the parameters, *c*_50_ (top) and *R*_*max*_ (bottom), for unmasked (left) and masked (right) responses from 1F1 (A) and 2F1 (B) SSVEP components. Notches and whiskers indicate the 95% confidence interval and range of bootstrap estimates respectively. Significant differences in comparison to control flies are indicated by asterisks: * = *p* <.050; ** = *p* <.010, *** = *p* <.001. **(C, D)** Hyperbolic functions modelling CRFs based on the means of bootstrapped parameters for 1F1 (C) and 2F1 (D) components for unmasked (top) and masked (bottom) responses.

